# Heading direction tracks internally directed selective attention in visual working memory

**DOI:** 10.1101/2022.05.04.490654

**Authors:** Jude L. Thom, Anna C. Nobre, Freek van Ede, Dejan Draschkow

**Author notes:** These authors contributed equally (Freek van Ede, Dejan Draschkow). Corresponding authors: Jude Thom, Dejan Draschkow.

## Abstract

We shift our gaze even when we orient attention internally to visual representations in working memory. Here, we show the bodily orienting response associated with internal selective attention is widespread as it also includes the head. In three virtual reality (VR) experiments, participants remembered two visual items. After a working memory delay, a central colour cue indicated which item needed to be reproduced from memory. After the cue, head movements became biased in the direction of the memorised location of the cued memory item – despite there being no items to orient towards in the external environment. The heading-direction bias had a distinct temporal profile from the gaze bias. Our findings reveal that directing attention within the spatial layout of visual working memory bears a strong relation to the overt head orienting response we engage when directing attention to sensory information in the external environment. The heading-direction bias further demonstrates common neural circuitry is engaged during external and internal orienting of attention.

## Introduction

We often move our head when orienting attention overtly to sensory information in our environment. We can also orient attention covertly to items in the external world, in the absence of large head movements. For example, you may be watching a film whilst directing your attention towards your phone when you are expecting a phone call. Covertly orienting attention to items in the environment is accompanied by subtle overt manifestations of orienting behaviour, including directional biases in eye-movements (Engbert & Kliegl, 2003; Hafed et al., 2011; Hafed & Clark, 2002; Yuval-Greenberg et al., 2014).

We can also orient attention internally to items maintained in the spatial layout of visual working memory (Griffin & Nobre, 2003; Manohar et al., 2019; Murray et al., 2013; Olivers et al., 2011; Souza & Oberauer, 2016; van Ede & Nobre, 2021). Similar to attentional selection in the external world, internal selective attention within visual working memory is associated with small directional eye-movement biases toward the memorised locations of attended items (Draschkow et al., 2022; van Ede, Chekroud, & Nobre, 2019; van Ede et al., 2020, 2021) (see also: (Ferreira et al., 2008; Spivey & Geng, 2001)). This overt manifestation of internal selective attention occurs despite the external absence of the attended memory items and even when memorised item location is not required for task performance.

Head movements are also affected by covert attentional selection. Covert attention activates neck muscles (Corneil et al., 2007; Corneil & Munoz, 2014a) and the lag between head and eye movements is affected by the congruency of covert attentional cues (Khan et al., 2009), suggesting that the head and eyes may each be modulated or involved when directing covert attention towards items in the external environment. The potential involvement of head and eye movements may be separable, provided there are differences in the neurophysiological pathways controlling head and eye movements (Gandhi & Sparks, 2007). Therefore, it is important to explore both the head and eyes when asking questions relating to bodily orienting behaviour, since the head and eyes may contribute in distinct ways as part of a broader bodily orienting response (Corneil & Munoz, 2014b).

If the overt ocular traces of covert selective attention in memory (Draschkow et al., 2022; van Ede, Chekroud, & Nobre, 2019; van Ede et al., 2020, 2021) are part of a more widespread bodily orienting response, then directing internal selective attention to items in working memory should also be accompanied by head movement. Therefore, it is conceivable that internally directed selective attention in working memory may not only be associated with small orienting behaviour of the eyes but also of the head.

To test whether such an embodied orienting response of eyes and head occurs during internally directed spatial attention, we analysed head- and eye-tracking data from a VR study investigating selective attention in visual working memory (Draschkow et al., 2022). The head-tracking data, which was not interrogated previously, allowed us to address whether head movements are similarly biased towards the memorised locations of selectively attended items in visual working memory.

## Materials and Methods

The data were collected as part of a study which used VR to examine different spatial frames of working memory in immersive environments (Draschkow et al., 2022). To answer the current research question, we focused on head-movement data, which were not analysed in the previous study (Draschkow et al., 2022). In this section, we describe the experimental materials and methods relevant to the focus of our research question. Information on additional manipulations which were not the focus of the current study can be found in (Draschkow et al., 2022).

### Participants

We analysed data from three experiments (1-3). Each experiment had a sample size of 24 human volunteers. Sample size was based on our prior study that contained four experiments using a similar outcome measure (van Ede, Chekroud, & Nobre, 2019) and revealed robust results with 20–25 participants. To address our new research question and further increase power and sensitivity we combined the samples from the individual experiments to create a larger data set with 48 participants and 72 experimental runs. The participants in Experiments 1-2 were the same and were recruited separately from the participants in Experiment 3 (Experiments 1-2: mean age 25.8 years, age range 18-40 years, all right-handed, 20 female; Experiment 3: mean age 25.5 years, age range 19-37 years, 1 left-handed, 13 female). All participants had normal or corrected-to-normal vision. Participants provided written consent prior to the experiments and were compensated £10 per hour. Protocols were approved by the local ethics committee (Central University Research Ethics Committee #R64089/RE001 and #R70562/RE001).

### Materials and Apparatus

Participants wore an HTC Vive Tobii Pro VR headset. Participants held the controller in their dominant hand, using their index finger and thumb to press response buttons. The positions of the headset and hand controller were recorded by two Lighthouse base stations, using 60 infrared pulses per second. These pulses interacted with 37 sensors on the headset and 24 sensors on the controller, providing sub-millimetre tracking accuracy. The headset contained a gyroscope and accelerometer, allowing for the precise recording of head rotational positions (accuracy < 0.001°). The headset contained a binocular eye tracker (approximately 0.5° visual angle accuracy, sampling rate 90 Hz). Two OLED screens displayed the environment in the headset (refresh rate 90 Hz, 1080 x 1200 pixels, field-of-view 100° horizontal × 110° vertical). We used Vizard (Version 6) to render and run the VR experimental environment on a Windows desktop computer.

In the VR environment, participants stood in the centre of a virtual room (4.2 m long, 4.2 m wide, 2.5 m tall) with a grey concrete texture applied to the four walls (Fig 1A). The working memory items were two coloured bars (length 0.5 m/14.25° visual angle, diameter 0.05 m /1.425°of visual angle), which appeared 2 m in front of the participant. One item appeared 1 m to the left (28.7° visual angle), on the front wall. The other appeared 1 m to the right, on the front wall. The centres of the items were 2 m apart.

**Fig 1.**
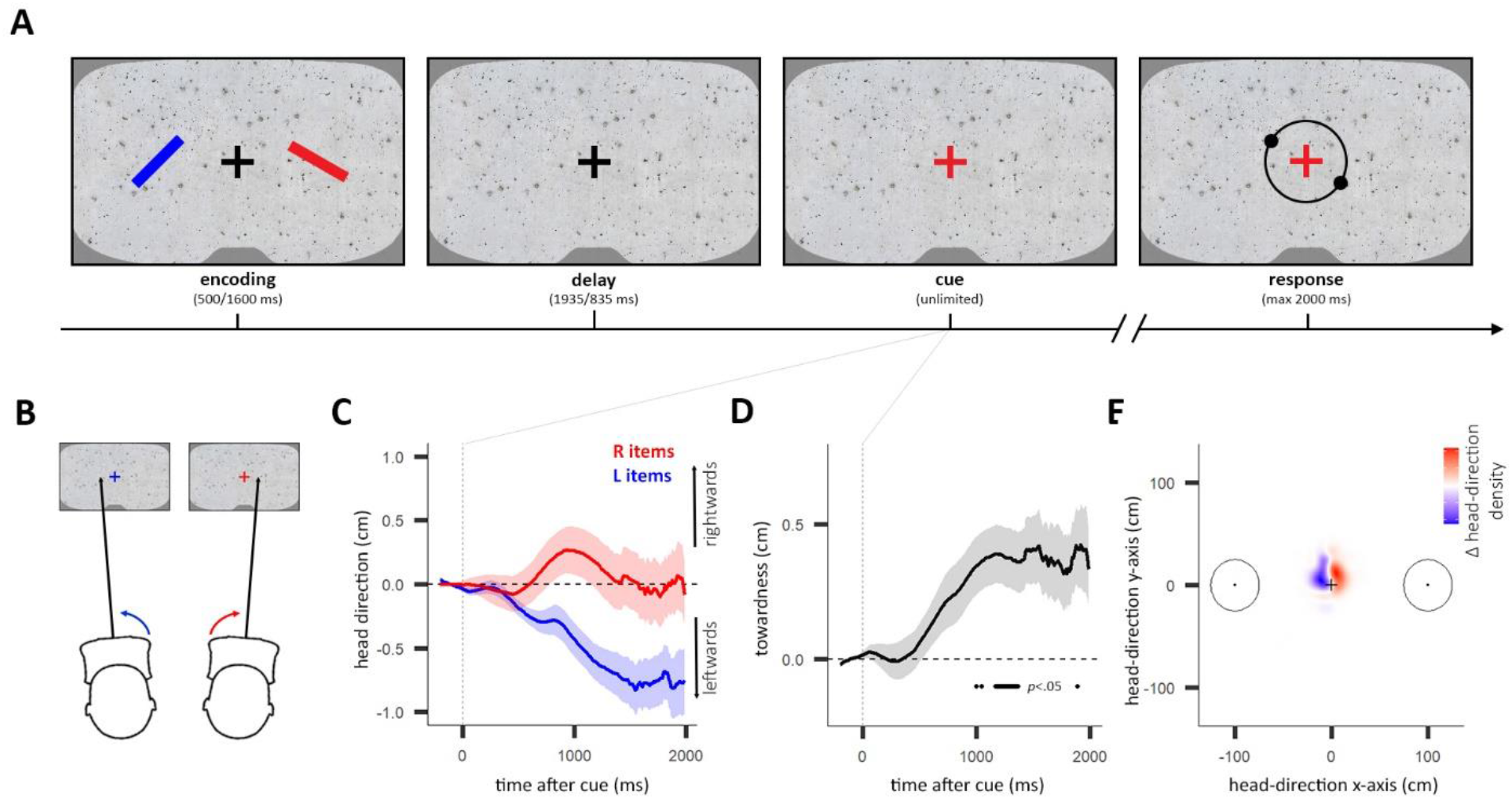
Heading direction tracks attentional selection in visual working memory. **A)** Participants remembered the orientations of two coloured items in a VR environment. After a delay, the fixation cross retrospectively cued the colour of the target item. Participants then reported the orientation of the target item using the controller. **B)** We recorded the shift in the projected location (in cm) of the “heading direction” onto the virtual wall in front of the participant. **C)** Average heading direction for left (L) and right (R) item trials as a function of time after cue. Shading indicates ± 1 SEM. **D)** Towardness of heading direction as a function of time after cue. Horizontal line indicates a significant difference from zero, using cluster-based permutation testing. Shading indicates ± 1 SEM. **E)** Density map showing the difference in heading-direction density between right minus left item trials (500-2000 ms after cue). Circles indicate the locations of the items during encoding. Centres of items are at 100 cm (28.7 degrees of visual angle) **C-E)** Data aggregated from Exp 1-3. See Fig A1 for separate plots of heading direction and heading-direction towardness as functions of time after cue for Exp 1-3.

### Procedure and Tasks

Participants were given time to get used to the headset, controller, lab room, and virtual environment before the experiments began. This included 24 practice trials in which participants learnt how to make responses and became familiar with the trial sequence.

In all experiments, each trial consisted of the same main steps (Fig 1A). At the beginning of each trial, participants stood upright in the centre of the room and were instructed to fixate on a fixation cross with their eyes (size 12 cm x 12 cm, ~3.4° visual angle). During the task, participants were free to hold their heads as they liked. After 500 ms of fixation, two items appeared (as described in Materials and Apparatus). Both items were slanted at independently drawn random orientations (ranging 0-180°). One item was red, and the other was blue. The colour of each item was allocated randomly on each trial. Participants were instructed to remember the orientations of the items during a delay.

All three experiments included conditions in which participants turned 90° to the left or right during the delay between the presentation of the items and the cue (“turning trials”). These turning trials were part of a separate study addressing a distinct question regarding how selection dynamics in visual working memory are influenced by self-movement (Draschkow et al., 2022) and were not included in our analyses.

Due to differences in the turning trials between experiments, the timings of the tasks differed between experiments. In Experiment 1, the items disappeared after 500 ms, compared to Experiments 2-3 where the items remained present for 1600 ms. After the items disappeared, the participant needed to remember the orientations of the items during a delay. The delays lasted 1935 ms (Experiment 1) and 835 ms (Experiments 2-3) after the items disappeared.

Following the delay, the fixation cross changed to a blue or red colour – matching the colour of the left or right item in working memory. The colour cue indicated the item for which the orientation response needed to be reproduced (target item) and signalled that participants could initiate the response when ready. The target item was randomly selected in each trial irrespective of orientation, location, and colour. Participants had unlimited time to recall the orientation of the target item and activate a response.

Once a response was initiated, participants had 2000 ms to dial in the orientation of the target item, using the controller. The response activation generated a dial made of two handles (diameter 0.06 m) on a circular torus (diameter 0.5 m, tube diameter 0.02 m), which was centred at the fixation cross. This dial was only present during the response stage. The handles moved along the torus according to the controller’s orientation, allowing participants to reproduce the orientation of the target item. Participants confirmed their response by pressing the trigger button of the controller. Immediately after confirming their response, the dial disappeared, and participants received feedback on their performance. Feedback was presented on a 0-100 scale, with 100 being perfect reproduction of the target item’s orientation. This number was presented above the fixation cross for 500 ms. Feedback was followed by a 700 ms delay. After this delay there was an inter-trial interval randomly selected between 1500 and 2000 ms.

There were 100 stationary trials in each experiment (50 left target item, 50 right target item). Trials were presented in 5 blocks with 20 trials each. The headset re-calibrated the gaze tracking at the beginning of each block. Participants completing Experiments 1 and 2 performed both tasks in the same session, in counterbalanced order. Each experiment lasted approximately one hour, and the full session lasted approximately two hours.

### Data Analysis

Tracking and behavioural recordings were stored in a comma-separated variable file (CSV), for each participant. We used R Studio (Version 1.3.1093, 2020) to analyse the data. The data files and analysis scripts are available online here: *https://doi.org/10.17605/OSF.IO/24U9M*

The “heading direction” variable refers to the projected location (in cm) of the heading direction onto the virtual wall in front of the participant. The “gaze direction” variable was the horizontal distance between the fixation cross and the gaze-fixation point on the virtual wall (averaged between both eyes). For an illustration of the heading direction variable, see Fig 1B.

We also recorded yaw, roll, and translation of the headset (Fig 2) to look at the contributions of these individual components of the heading-direction vector. Head yaw is the rotational position around the head’s vertical axis. Head roll is the rotational position around the head’s longitudinal axis. For example, rotating your head whilst reading a large sign left-to-right would be reflected in changing yaw values and tilting your head to read a slate sign would change roll values. Translation refers to the horizontal movement of the entire headset (e.g., if the participant moved their entire head to the left whilst looking straight ahead). Together, yaw, roll, and translation are components which can influence the horizontal heading direction.

**Fig 2.**
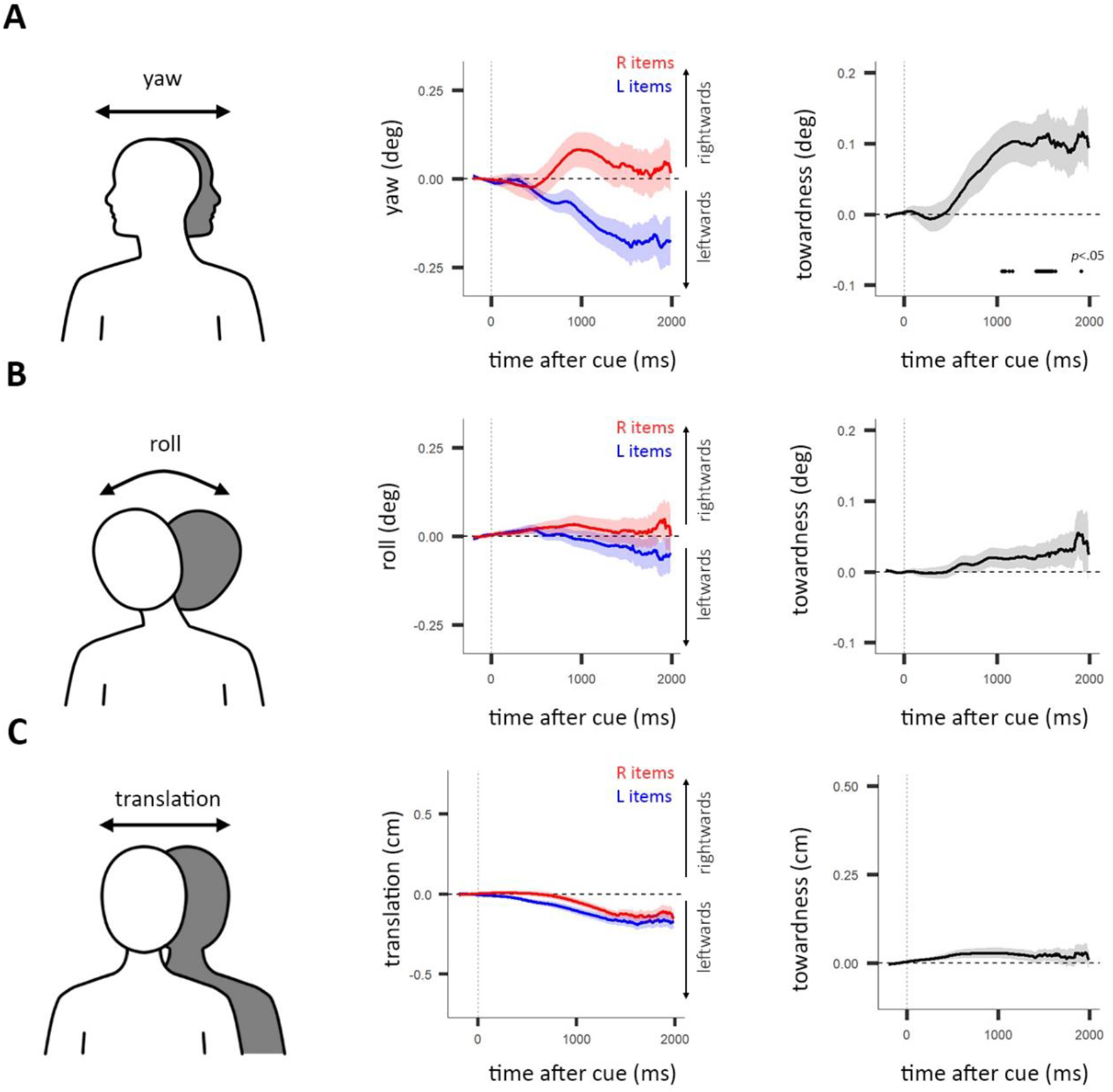
Biased movement in yaw, roll, and translation. **A)** Left panel) Yaw as a component of heading direction. Centre panel) Average yaw for left (L) and right (R) item trials as a function of time after cue. Shading indicates ± 1 SEM. Right panel) Towardness of yaw as a function of time after cue. Horizontal line indicates a significant difference from zero, using cluster-based permutation testing. Shading indicates ± 1 SEM. **B)** Same as A, using roll instead of yaw. **C)** Same as A, using translation instead of yaw. **B-C)** Lack of horizontal lines in towardness plots in the right panels indicates no significant difference from zero was found, using cluster-based permutation testing with a threshold of *p*<.05.

We epoched the data from 500 ms before cue to 2000 ms after cue. We smoothed all time-course data over 4 samples (44 ms smoothing window average). In each trial, the mean value between 200 and 0 ms before the cue was used as a baseline and subtracted from all values in the trial. We excluded trials in which heading direction or gaze direction exceeded 0.5 m (half the distance to the locations of the memoranda) in either direction of the fixation cross during the time window (−500 ms – 2000 ms) to remove the effect of large outliers. This cut-off was set a priori in accordance with our previous work (Draschkow et al., 2022). We also excluded trials with a yaw or roll of over 20 degrees in either direction (average percentage of excluded trials per participant: M 5.96%, SE 0.01; total percentage of excluded trials: 16.58%). Importantly, however, not applying any cut-off did not change the findings presented in the Results section.

We compared behaviour between right- and left-item trials in the three experiments separately to check if the side of the target item affected performance. We used within-subject ANOVAs to check for effects of target side on error and response time. To follow-up findings (including null findings), we conducted Bayesian t-tests (Rouder et al., 2009) with the default settings of the ‘Bayes Factor’ package (Richard D. Morey et al., 2021). Bayes-factor values either indicated evidence in favour of the alternative hypothesis (B_01_ > 3), in favour of the null hypothesis (B_01_ < 0.33) or suggested inconclusive evidence (B_01_ > 0.3 and B_01_ < 3)(Kass & Raftery, 1995).

Next, we plotted the change in the time-course data (heading direction, yaw, roll, translation, gaze direction) from baseline (−200 – 0 ms before cue), separately for left- and right-item trials. To increase sensitivity and interpretability, we constructed a single measure of ‘towardness’. Towardness aggregated horizontal movement towards the target item on each trial, combining leftward movement in left-item trials and rightward movement in right-item trials. A positive towardness indicated a horizontal position in the direction of the target item. Towardness for each time step was given by the trial-average horizontal position in right-item trials minus the trial-average horizontal position in left-item trials (where position values left of fixation were negative) divided by two. The same procedure for calculating towardness was used for all time-course head and gaze data. We used this towardness variable to determine the significance of the biased movements (compared to zero), using “cluster-depth” (Frossard & Renaud, 2022) cluster-based permutation tests (Maris & Oostenveld, 2007; Sassenhagen & Draschkow, 2019). We ran the cluster-based permutation testing in R with the ‘permuco’ package (Frossard & Renaud, 2022; Frossard & Reneud, 2021).

To gain a better understanding of the scale and variance of the heading direction, we plotted a density map of all of the heading-direction values between 500 ms and 2000 ms post-cue over all trials and all participants (including excluded trials). We used colour to code the side of the target item in the trial and highlight differences in the directionality of heading direction between item-sides.

## Results

Participants performed a visual working memory task in a VR environment while we tracked their head and gaze. In the task, participants remembered the orientations of two coloured bars, one on the left and one on the right, for a short delay (Fig 1A). After the working memory delay, a colour cue indicated the bar for which participants needed to reproduce the orientation on a dial.

### Heading direction tracks internal selective attention in visual working memory

After the colour change in the fixation cross (cue onset), horizontal heading direction became biased in the direction of the memorised external location of the cued memory item (Figs 1B-1E). This heading-direction bias occurred although there was no information present or expected at the external location corresponding to the memorized item after the colour cue.

The bias in horizontal heading movement was leftwards in trials in which the colour cue corresponded with the memory item that had been encoded on the left (“left item”), and rightwards in trials in which the colour cue corresponded with the memory item that had been encoded on the right (“right item”). Fig 1B illustrates the nature of the heading-direction bias in left- and right-item trials. The average heading direction after the colour cue for trials with cued memory items on the left and right are plotted separately in Fig 1C. To quantify this heading-direction bias and express it as a single measure, we combined the heading-direction bias from left and right item trials into a measure of towardness (van Ede, Chekroud, & Nobre, 2019). The towardness of the heading direction became evident starting at approximately 500ms after the onset of the cue (Fig 1D; cluster p < .05; largest cluster ranging between 1167-1367 ms).

To explore the scale of the heading-direction bias, we calculated density maps of single-trial heading-direction values, and subtracted density maps between left-and right-item trials. To focus on the window of interest, we considered all heading-direction values when the heading-direction bias was most pronounced (500-2000 ms; Fig 1E). This revealed the subtle nature of the heading-direction bias. Participants did not move their heading direction all the way to the memorised locations of the items (circles in Fig 1E). Instead, participants subtly moved their heading direction towards the memorised item locations (< 0.5° of rotation), with heading-direction biases remaining close to fixation – akin to the type of directional biases we have recently observed in gaze (Draschkow et al., 2022; van Ede, Chekroud, & Nobre, 2019; van Ede et al., 2020, 2021). The properties of the heading-direction bias were similar across three slightly different versions of the task (Exp 1-3) and are plotted separately in Fig A1. There were no significant effects of target side (left vs right) on behavioural performance (error and response time) in any of the experiments (see Fig A2).

### The heading-direction bias is driven by movement along the head’s yaw axis

To determine which heading-movement components contributed to the heading-direction bias, we separately analysed yaw, roll, and translation. Like the heading-direction vector, yaw followed the movement pattern of heading-direction in left- and right-item trials, which was also confirmed by a significantly positive towardness cluster (Fig 2A; p < .05, cluster-corrected). Roll showed a non-significant towardness trend (Fig 2B; p > 0.999 for all clusters of the full time window) and translation did not move towards the memorised locations of the cued memory items (Fig 2C; p > 0.257 for all clusters of the full time window). We also investigated all components making up the heading direction measure (x-, y-, z-translation, yaw, pitch, and roll), during the critical 500-1500 ms post-cue period (Fig A3). Fig 3 shows how head rotation around the yaw-axis closely tracks heading direction. Thus, the leftwards-rightwards rotation along the head’s yaw axis was the primary factor contributing to the directional heading-direction bias when selectively attending items in our visual working memory task.

**Fig 3.**
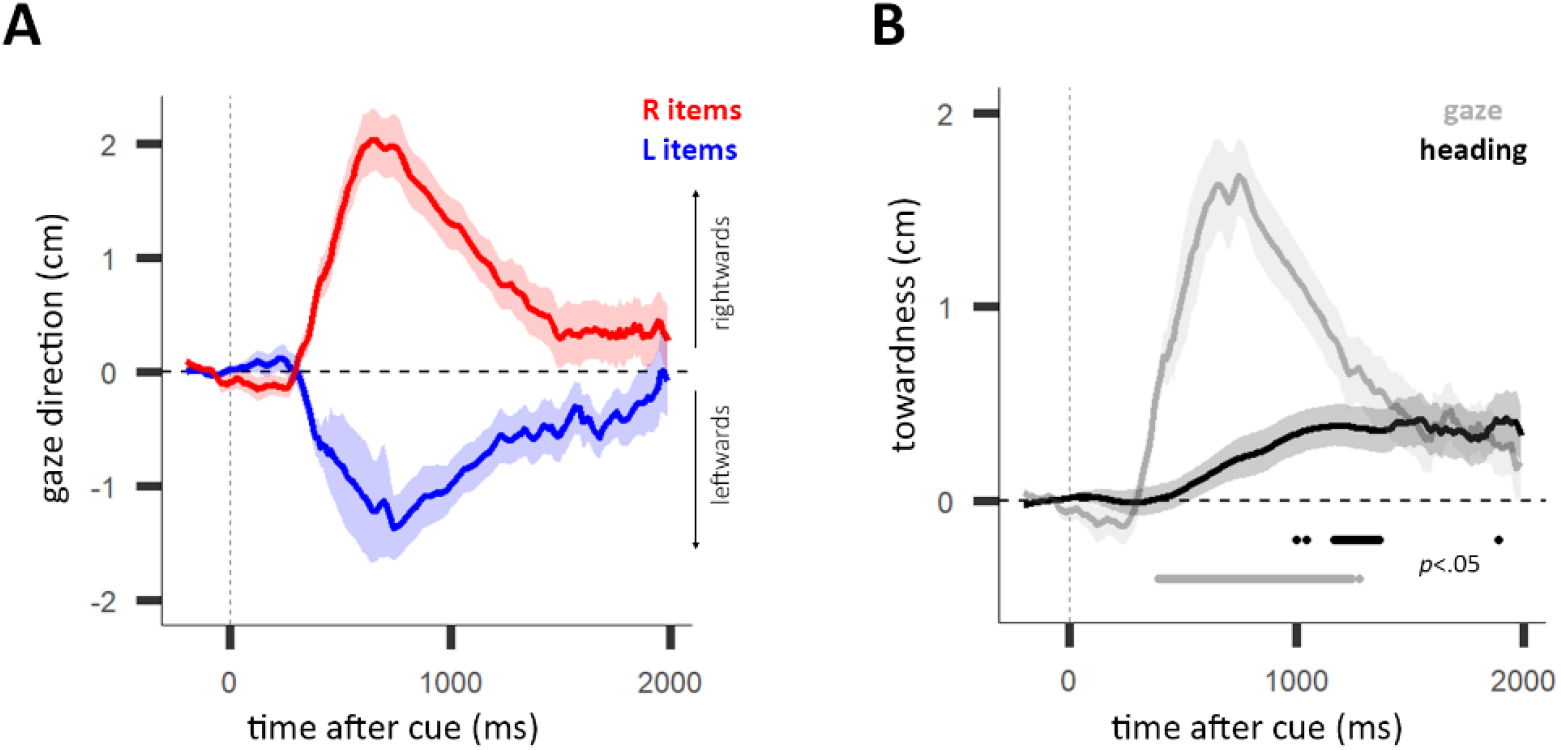
The gaze bias and the heading-direction bias. **A)** Average gaze direction for left (L) and right (R) item trials as a function of time after cue. Shading indicates ± 1 SEM. **B)** Towardness of gaze direction (gaze) and heading direction (heading) as a function of time after cue. Horizontal line indicates a significant difference from zero, using cluster-based permutation testing. Shading indicates ± 1 SEM.

### The heading-direction bias is accompanied by a gaze bias in the same direction

Like the heading direction, gaze direction moved towards the location of the cued item during internal selective attention (as we have previously reported in this dataset (Draschkow et al., 2022) as well as in prior datasets (van Ede, Chekroud, & Nobre, 2019; van Ede et al., 2020, 2021; van Ede & Nobre, 2021)). Fig 3A shows the leftwards and rightwards movement of gaze direction in left and right item trials. The gaze towardness was significantly different from zero after the cue (p < .05 between 400-1244 ms, cluster-corrected; Fig 3B). We focused our statistical analyses on the data aggregated across the individual experiments to improve sensitivity, noting that the gaze direction for left and right trials and towardness over time were similar between all three experiments (Exp 1-3; Fig A1).

In Fig 3B, we also overlay the heading-direction bias for a descriptive comparison. While the heading-direction bias and gaze bias both acted toward the memorised location of the cued item, Fig 3B shows the bias lags behind the gaze bias. The largest significant cluster (Frossard & Renaud, 2022; Frossard & Reneud, 2021) for the gaze bias was significant at ~400ms, whereas the significant time window for the heading-direction bias started more than a full second after the cue (p < .05; heading: 1167-1367 ms, gaze: 400-1244 ms).

## Discussion

Our results reveal that, like eyes, the heading direction tracks internally directed selective attention inside visual working memory. This manifests in directionally biased head movements towards the memorised location of attended memory items. Although the heading direction bias is small (Fig 1 & Fig A1), we were able to capture it by calculating the relative change in heading direction triggered by the cue and by aggregating the data from multiple experiments. The heading-direction bias in our task was predominantly driven by the head’s rotation around its yaw-axis and accompanies a gaze bias in the same direction. The observed heading-direction bias suggests there is a general bodily orienting response during internal selective attention – suggesting brain structures involved in orienting of the eye and head are also engaged when orienting within the internal space of working memory.

The heading-direction and gaze biases may reflect bodily signatures which are part of a widespread orienting response activating brain areas that are involved in both overt and covert attentional selection. Indeed, there is good evidence that the brain’s oculomotor system is also involved in covert orienting of spatial attention (Deubel & Schneider, 1996; Engbert & Kliegl, 2003; Hafed et al., 2011; Hafed & Clark, 2002; Moore & Armstrong, 2003; Moore & Fallah, 2004; Nobre et al., 1997; Yuval-Greenberg et al., 2014). Moreover, from an evolutionary perspective, it is conceivable that our ability to orient internally evolved gradually from external orienting behaviours of the head and eyes – maybe relying on overlapping neural circuitry (Cisek, 2019). From this perspective, the observed subtle bias in head- and eye-movements may reflect an inevitable ‘spill over’ from activating neural circuitry that has evolved to orient both internally and externally (Strauss et al., 2020).

It is maybe surprising to find this heading-direction bias, even when attention is directed internally and without any items in the environment toward which to orient. However, in natural settings, there may be a behavioural benefit of orienting the head and eyes towards the locations of selected memory items. In our task, no subsequent behavioural goal benefited from orienting towards the memorised location of the attended memory item. However, in daily life, items rarely disappear from their location in the external environment as they do in our task. Thus, orienting the eyes and head towards the memorised locations of selected items may serve to guide future behaviour, such as resampling items. In fact, people often resample items in a naturalistic working memory task, when it is easy to do so (Ballard et al., 1995; Draschkow et al., 2021). For example, imagine you are with a friend in a café, and they comment on the barista’s hat. You may attend the barista in memory, attempting to recall what their hat looked like. At the same time your head and eyes may be preparing for you to shift your gaze and look at the barista’s hat again. In this way, the small heading-direction and gaze biases towards selected items in working memory may reflect a natural tendency to engage in action in relation to selected memoranda (Boettcher et al., 2021; Heuer et al., 2020; Olivers & Roelfsema, 2020; van Ede, 2020; van Ede, Chekroud, Stokes, et al., 2019), even if there was no incentive for this in our task.

In natural behaviour, head and eye movements are intrinsically functionally linked (Foulsham et al., 2011; Land, 2009; Solman et al., 2016) and head movements can even compensate for eye movements when people cannot make saccades (Ceylan et al., 2000; Gilchrist et al., 1997). This coordinated relationship between head- and eye-movements motivated us to look at both the head and eyes when exploring bodily orienting responses. The heading-direction bias revealed here implicates that neural circuitry which controls head movements – at least along the yaw axis – is recruited by, and potentially overlaps with, circuitry that directs internal selective attention. In fact, previous research has found overlap between brain areas thought to process spatial attention and eye and head movements. For example, the frontal eye fields (FEF) play a role in directing attention and controlling eye movements (Bruce & Goldberg, 1984; Grosbras & Paus, 2002; Moore & Fallah, 2004; Robinson & Fuchs, 1969; Taylor et al., 2007). Alongside attentional selection and eye movements, the FEF also contributes to head movements. The hemodynamic activity of the FEF responds to head movement (Petit & Beauchamp, 2003) and microstimulation to the FEF in primates results in head movement (Chen & Walton, 2005; Elsley et al., 2007). Additionally, modulation of activity in the superior colliculus – an area shown to process not only eye (Schiller & Stryker, 1972; R. Wurtz & Albano, 2003; R. H. Wurtz & Goldberg, 1971) but also head movements (Bizzi et al., 1972; Corneil et al., 2002) – also affects the deployment of covert attention (Krauzlis et al., 2013; Lovejoy & Krauzlis, 2009; Müller et al., 2005). Our results complement these findings, with the heading-direction and gaze biases suggesting overlap between neural circuitry and activity governing attentional selection inside working memory, eye movements, and head movements.

However, control of the head and eye is not entirely linked, as shown by differences in the neurophysiological pathways controlling eye and head movements (Bizzi et al., 1972; Horn et al., n.d.; Oommen & Stahl, 2005). This is demonstrated in the distinct temporal profiles of the heading-direction and gaze biases presented here which highlight the value of looking at multiple components of what might be a widespread bodily orienting response involving the head and eyes. It is important to note that comparisons between the temporal profiles of the head and gaze biases should be made with caution due to differences in mass and musculature of the head and eyes and the signal-to-noise ratio of the two measures.

It is worth noting the apparent asymmetry in the magnitude and time course of the heading-direction bias in left vs right trials and across experiments (as seen in Fig 1 and Fig A1). Based on our previous work on gaze biases (Draschkow et al., 2022; van Ede, Chekroud, & Nobre, 2019; van Ede et al., 2020, 2021), we a-priori decided to focus on a single measure of ‘towardness’, which represents horizontal movement towards the target item on each trial. This aggregated measure does not only benefit from increased sensitivity, but also removes any potential drifts in the measure that are not due to selective attention (that could potentially contribute to the apparent asymmetry we observed here). In future studies it would be interesting to further investigate these potential asymmetries, and how they relate to behavioural performance, for example by increasing trial numbers and introducing a neutral condition in which no item is cued.

Finally, by using VR, we were able to measure the heading-direction bias alongside the gaze bias as participants’ head, eye, and body were unconstrained. To date, the benefits of VR have been appreciated most prominently by researchers studying naturalistic human navigation, ethology, and long-term memory (Draschkow & Võ, 2017; Helbing et al., 2020; Li et al., 2016; Mobbs et al., 2021; Stangl et al., 2020; Topalovic et al., 2020). Our present findings further highlight the benefits of using VR (combined with eye- and head-tracking) to study bodily orienting behaviour (Draschkow et al., 2021, 2022) related to internal cognitive processes, as showcased here for internal attentional focusing in working memory.

## Data Availability Statement

The data files and analysis scripts are available online here: https://doi.org/10.17605/OSF.IO/24U9M

## Author Contribution

J.L.T. - Formal Analysis, Investigation, Visualization, Writing – original draft, Writing – review & editing. A.C.N. - Funding acquisition, Project administration, Resources, Supervision, Writing – original draft, Writing – review & editing. F.v.E - Funding acquisition, Investigation, Methodology, Project administration, Resources, Supervision, Writing – original draft, Writing – review & editing. D.D. - Data curation, Formal Analysis, Investigation, Methodology, Project administration, Resources, Supervision, Writing – original draft, Writing – review & editing.

## Appendices

**Fig A1.**
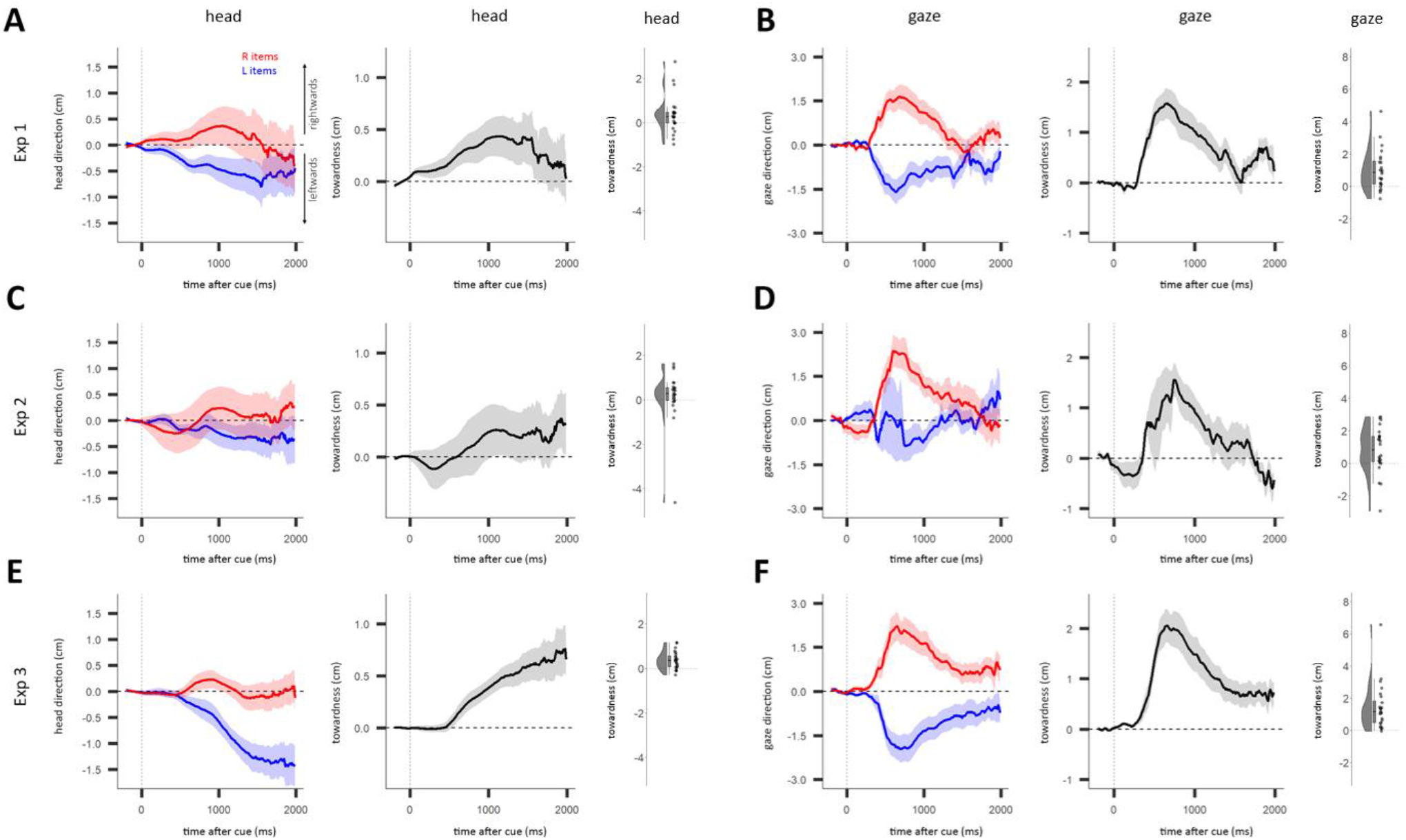
The heading-direction and gaze biases for Experiments 1-3. **A)** Left panel) Average heading direction for left (L) and right (R) item trials as a function of time after cue, using data from Exp 1. Middle panel) Towardness of heading direction as a function of time after cue, using data from Exp 1. Right panel) Distribution of the mean towardness between 500-1500 ms across participants. **B)** Same as A, using gaze direction instead of heading direction. **C)** Same as A, using data from Exp 2. **D)** Same as B, using data from Exp 2. **E)** Same as A, using data from Exp 3. **F)** Same as B, using data from Exp 3. **A-F)** Shading indicates ± 1 SEM.

**Fig A2.**
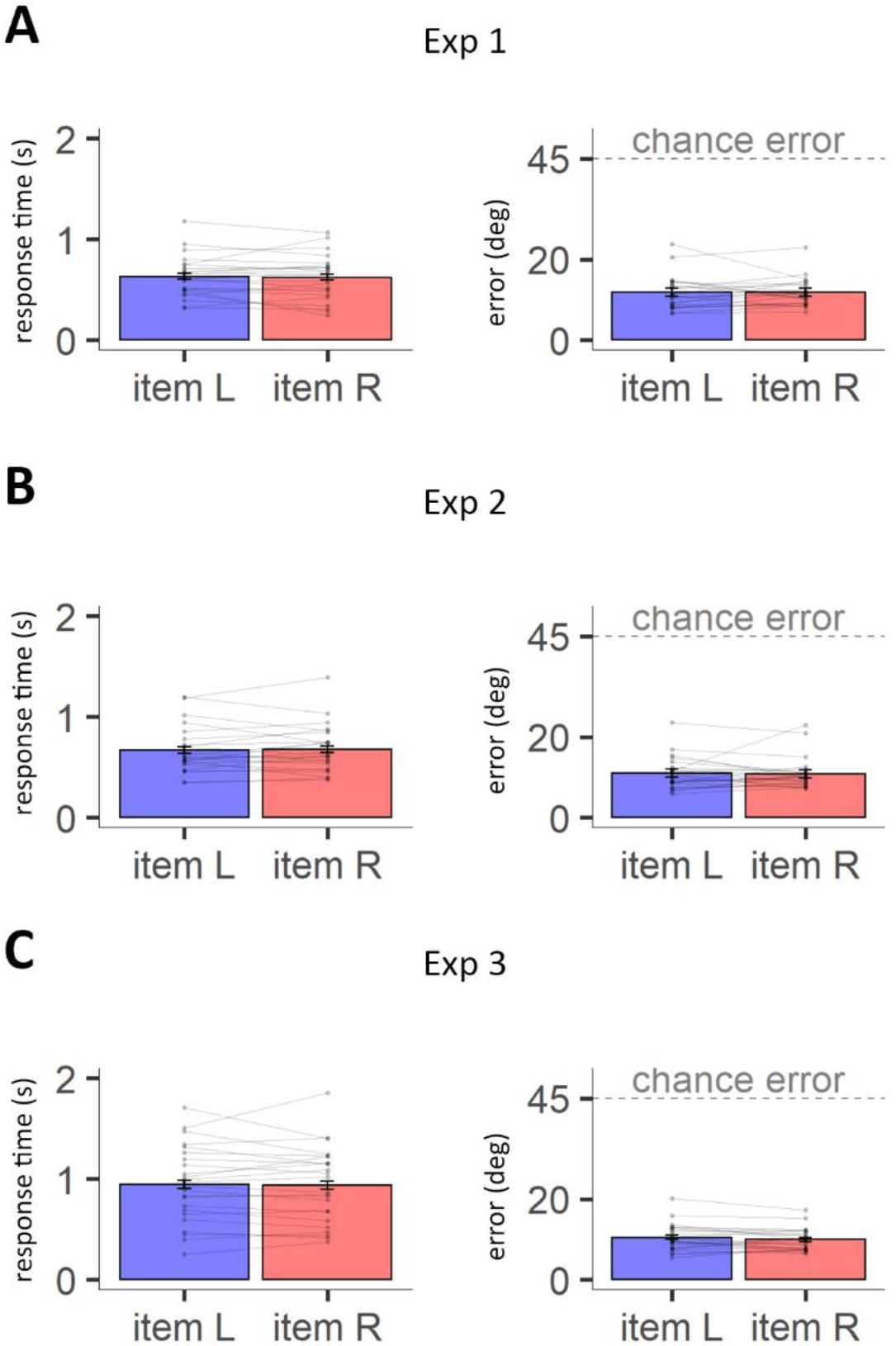
Similar performance in left and right item trials. **A)** Left panel) Plot comparing the mean response time between left item (item L) and right item (item R) trials, for each participant in Exp 1. Connected pairs of points are the means of the same participant. Error bars represent a 95% confidence interval. Right panel) Same as Left panel, for error instead of response time. **B)** Same as A, using data from Exp 2. **C)** Same as A, using data from Exp 3. There was no significant effect of target side on mean error in any of the experiments, Exp 1: (*F* (1,23) = 0.01, *p* = .934), Exp 2: (*F* (1,23) = 0.02, *p* = .881), Exp 3: (*F* (1,23) = 2.04, *p* = .166). For Exp 1-2, the follow-up Bayes t-test supported the null-hypothesis, suggesting the errors are similar between left and right item trials, Exp 1: (*B_01_* = 0.22), Exp 2: (*B_01_* = 0.22). Similarly, there was no significant effect of target side on mean response time in any of the experiments, Exp. 1: (*F* (1,23) = 0.19, *p* = .671), Exp 2: (*F* (1,23) = 0.23, *p* = .633), Exp 3: (*F* (1,23) = 0.07, *p* = .793). For Exp 1-3, the follow-up Bayes t-tests supported the null-hypothesis, suggesting the response times are similar between left and right item trials, Exp 1: (*B_01_* = 0.23), Exp 2: (*B_01_* = 0.24), Exp 3: (*B_01_* = 0.22).

**Fig A3.**
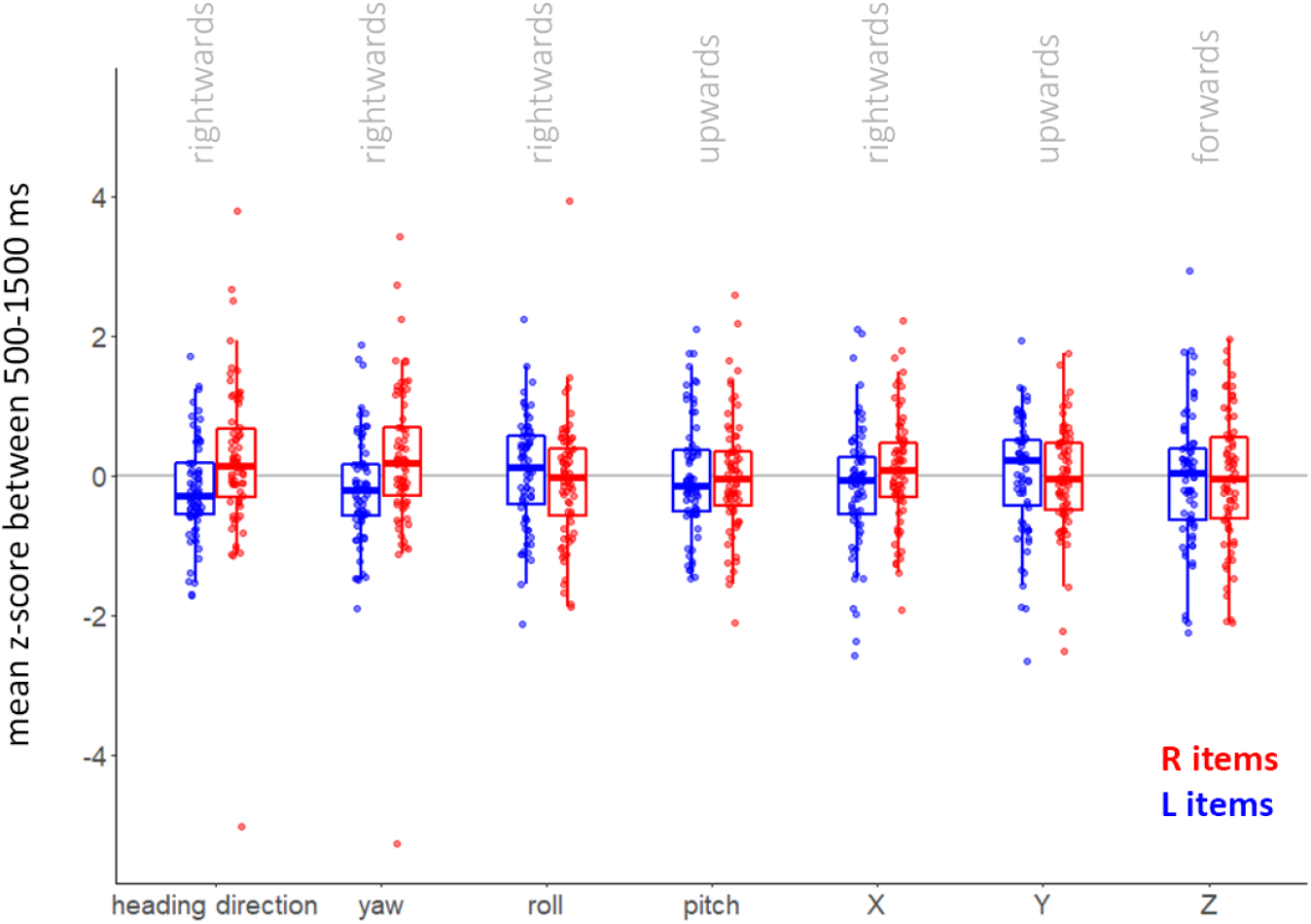
Distributions of measures making up heading direction. The measures which make up heading direction (x-, y-, z-translation, yaw, pitch, and roll) were z-score normalised before calculating their mean between 500-1500 ms. These mean values were averaged across blocks and trials for each participant and split by item side. Boxplots indicate median and interquartile range. The figure shows how head rotation around the yaw-axis closely tracks heading direction. We separately analysed yaw, roll, and X-translation in the main text and Fig

